# Mice tune out not in: Violation of prediction drives auditory saliency

**DOI:** 10.1101/633388

**Authors:** Meike M. Rogalla, Inga Rauser, Karsten Schulze, Lasse Osterhagen, K Jannis Hildebrandt

## Abstract

Successful navigation in complex acoustic scenes requires focusing on relevant sounds while ignoring irrelevant distractors. It has been argued that the ability to track stimulus statistics and generate predictions supports the choice what to attend and what to ignore. However, the role of these predictions about future auditory events in drafting decisions remains elusive. While most psychophysical studies in humans indicate that expected stimuli serve as implicit cues attracting attention, most work studying physiological auditory processing in animals highlights the detection of unexpected, surprising stimuli. Here, we tested whether in the mouse, target probability is used as an implicit cue attracting attention or whether detection is biased towards low-probability deviants using an auditory detection task. We implemented a probabilistic choice model to investigate whether a possible dependence on stimulus statistics arises from short term serial correlations or from integration over longer periods. Our results demonstrate that target detectability in mice decreases with increasing probability, contrary to humans. We suggest that mice indeed track probability over a time scale of at least several minutes but do not use this information in the same way as humans do: instead of maximizing reward by focusing on high-probability targets, the saliency of a target is determined by surprise.

## Introduction

Animals using acoustical information for navigation are nearly continuously confronted with numerous sounds from different sources. To differentiate between non-associated sounds that are present simultaneously, relevant stimuli need to be detected while irrelevant ones should be ignored. For this process of differentiation, the ability of tracking stimulus statistics (if a stimulus occurs with high or low probability) is essential, sets expectations, and creates predictions about future auditory events (Carbajal and Malmierca, 2018; Skerrit-Davis and Elhilali, 2018). While there is general agreement on the principal importance of expectation in auditory perception, there are different ways in which these predictions may be used to guide decisions.

One the one hand, high-probability, expected and relevant signals may attract selective listening, thus the ability to group and separate sounds from different sources and selectively pick and monitor one in the presence of others. This ability forms an important part of selective attention and supports the analysis of complex auditory scenes (Bregman, 1994; Sussman et al., 2007; Woods and McDermott, 2015). It has long been established that adult humans listen selectively for an expected auditory stimulus in reward-based auditory listening tasks (Scharf et al., 1987). Humans internally monitor the probability of a stimulus and adapt their behavior according to the stimulus statistics (Bargones and Werner, 1994a; Gordon Z. Greenberg and Larkin, 1968). This form of selective auditory attention does not require awareness of the subject and is driven by unconscious expectations (Wolmetz and Elhilali, 2016). Within this framework, the improvement of detectability is based on the expectation as an implicit cue and serves as an internal reward-maximizing strategy that drives the attention towards the expectation (Girshick et al., 2011).

While most psychophysical studies indicate that expected stimuli serve as implicit cues attracting attention, most work studying the physiology of auditory processing highlights the detection of unexpected, surprising stimuli. Stimuli are more salient when presented rarely to the auditory system and thus might be easier to detect due to pre-attentive mechanisms (Carbajal and Malmierca, 2018; Parras et al., 2017; Tiitinen et al., 1994). Within this framework, the evaluation of stimulus statistics serves to detect novelty, emphasizing changes in the auditory scene rather than enabling tracking of task relevant information.

Novelty coding in the auditory system has successfully been interpreted within the framework of predictive coding (Heilbron and Chait, 2017). Within this framework, surprising stimuli are carried on to higher sensory areas as prediction error signals, updating believes for the interpretation of sensory input. Whether and how prediction error signals shape the *detectability* of a signal is less clear: On the one hand, larger prediction error signals could result in higher detectability of surprising signals (den Ouden et al., 2012). On the other hand, (probability-cued) selective attention is thought to assign higher weights to the attended signal and thereby make it more salient (Kanai et al., 2015; Kok et al., 2012).

Thus, tracking of stimulus probability influences auditory processing in two contrary ways: on the physiological level, low-probability sounds elicit maximal responses, but during listening tasks, relevant high-probability sounds appear to attract attention, improving their detectability. While physiological evidence for deviant detection spans all the way from animal models to humans (Heilbron and Chait, 2017; Khouri and Nelken, 2015), behavioral assessment of the effects of target probability is largely restricted to humans.

Although rodents serve as widely used animal models to study auditory phenomena, little is known about their ability to monitor stimulus probability and its involvement in selective auditory attention. One study using chinchillas could not reproduce human results for auditory selective attention (Yost and Shofner, 2009). However, it remains unclear whether this generalizes to other rodents. Also, it is unknown if the animals simply do not adapt their behavior according to stimulus statistics, or if they rather respond towards unexpected, surprising stimuli instead of high-probability stimuli, as suggest by physiological data.

Here, we asked how target probability influences auditory perception in mice, as revealed in detection paradigms. More specifically, we tested whether target probability is used as an implicit cue attracting attention or whether detection is biased towards low-probability deviants. To this end, we devised three different psychophysical tasks and tested three separate sets of mice. First, we used faint tones in noise of different frequencies and varied the probability of a given tone frequency between different sessions. This paradigm resembles those used to test for the ‘listening band phenomenon’, the most prominent example of probability-guided attention in the human literature (Scharf et al., 1987). Subsequently, we tested whether the probability-dependence generalizes to other detection tasks, namely streaming paradigms, in which a target has to be detected in one out of multiple streams. Here we separately tested for effects on the detection of both spectral and temporal stimulus dimensions. Finally, we present a probabilistic choice model to investigate whether the dependence on stimulus statistics arises from short term serial correlations or from integration over longer periods.

## Results

### Experiment 1: Tone in noise detection

When humans are asked to detect faint tones in a noise background, performance for high-probability targets is better than for those played with low probability, even if listeners are not consciously aware of the probabilities (Greenberg and Larkin, 1968). This is usually explained by focused attention on specific auditory filters, thereby listing selectively to a certain frequency range (Bargones and Werner, 1994). In our first experiment, we aimed to test whether mice are able to track target probabilities from session to session and display a preference for either high- or low-probability targets. We devised a behavioral paradigm (Fig. 1), in which mice were trained to indicate the detection of faint tones embedded in a noise background by leaving a small pedestal after the presentation of a target (Fig. 1A). A typical single session contained 60 targets and lasted ~30 minutes. In order to test the animals near their individual thresholds, we first tested a single frequency in each session, varying the level of the tones to determine the threshold (Fig. 1C, upper panel). In the next step, we presented tones with varying probability as targets in mixed sessions (lower panel in (Fig. 1C). We hypothesized that if mice displayed selective listening to high-probability tones they should (1) be better at tone detection in the single frequency session compared to the mixed session and (2) show better performance for the high-probability compared to the low probability stimulus within the mixed session.

**Fig. 1.**
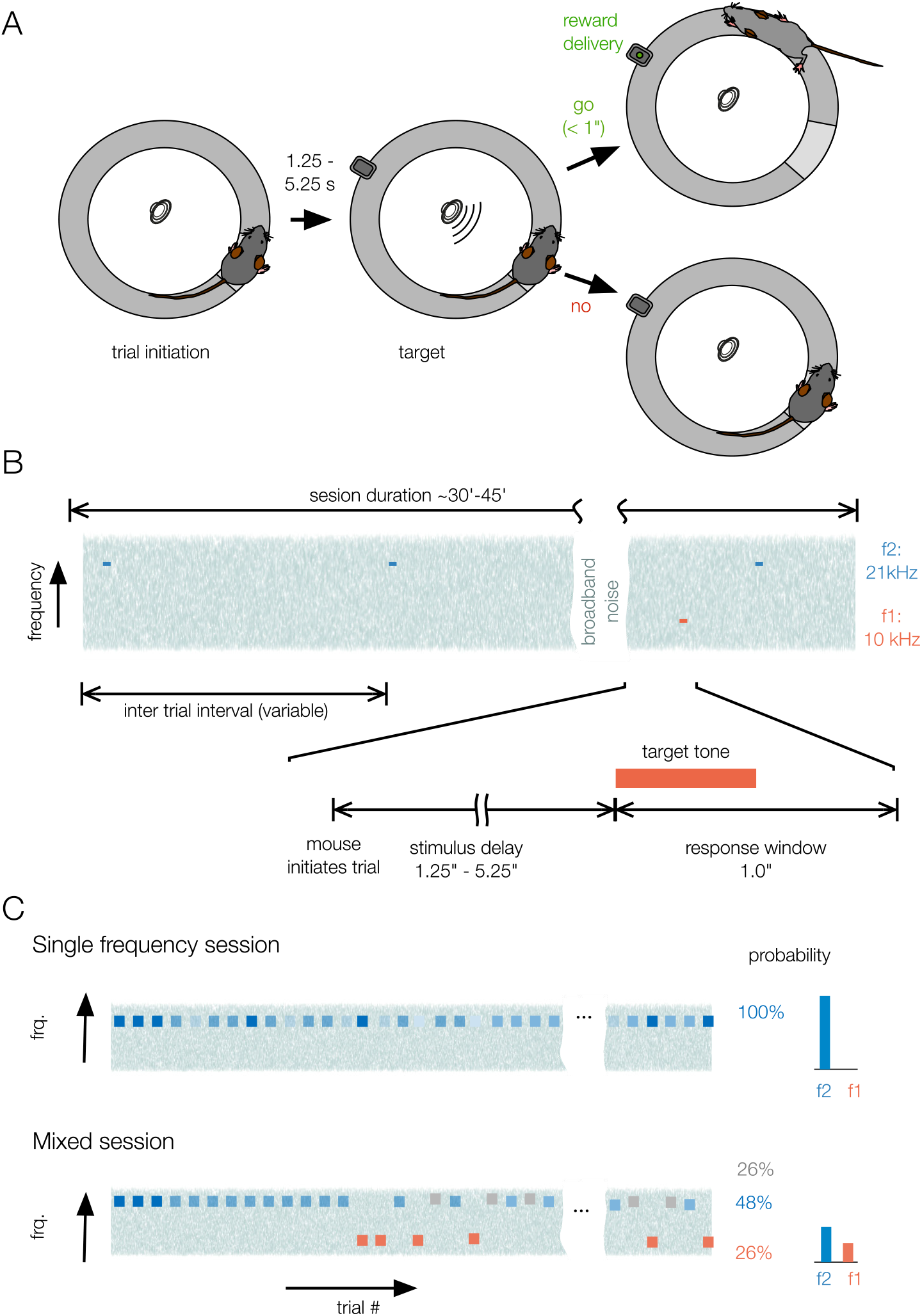
Behavioral paradigm and stimulus protocol used in Experiment 1. (A) Go/No Go paradigm used in this study. Mice initiated a trial by climbing on a small pedestal on the circular platform. After a variable waiting interval, a target was presented. Animals received a reward if they left the platform within 1s after target presentation. The next trial could be initiated immediately. (B) Timeline of one experimental session. Throughout the entire session, a broad band noise stimulus was presented. Once a trial was initiated, a 500ms pure tone was presented after a random stimulus delay. In a single session, an animal had to complete 73 or 78 trials, which typically lasted 30-45 minutes. (C) Different probabilities of single frequency pure tone targets in differents sessions. In single frequency sessions, the level of the tones was varied, but only pure tones of either frequency f1 (10kHz) or f2 (21kHz) were presented. In mixed session, level was held constant near the behavioral threshold, but three different frequencies were presented. In any one session, either f1 or f2 was presented with 48% probability and the respective other with only 26%. In addition, a tone of the frequency close to the high-probability targets was presented in 26% of the trials.

Contrary to our hypothesis, all animals tested showed higher sensitivity in the mixed than in the single frequency session tested before (example data in Fig. 2A; repeated measures ANOVA, F(1;20)=32.2, p<0.001). Within the mixed session, the impact of stimulus probability on the preference of the mice for low-probability tones was confirmed. Sensitivity was positively influenced by surprise, quantified as the prediction error (inverse conditional probability of the stimulus, Gill et al., 2008, Fig. 2B). This relation was highly significant, both when taking the single frequency sessions into account and for mixed sessions only, and independent of the frequency that was played. We concluded that mice are able to track target probabilities over a time frame of minutes to hours, but instead of the high-probability sound the low-probability sounds were detected more reliably.

**Figure 2.**
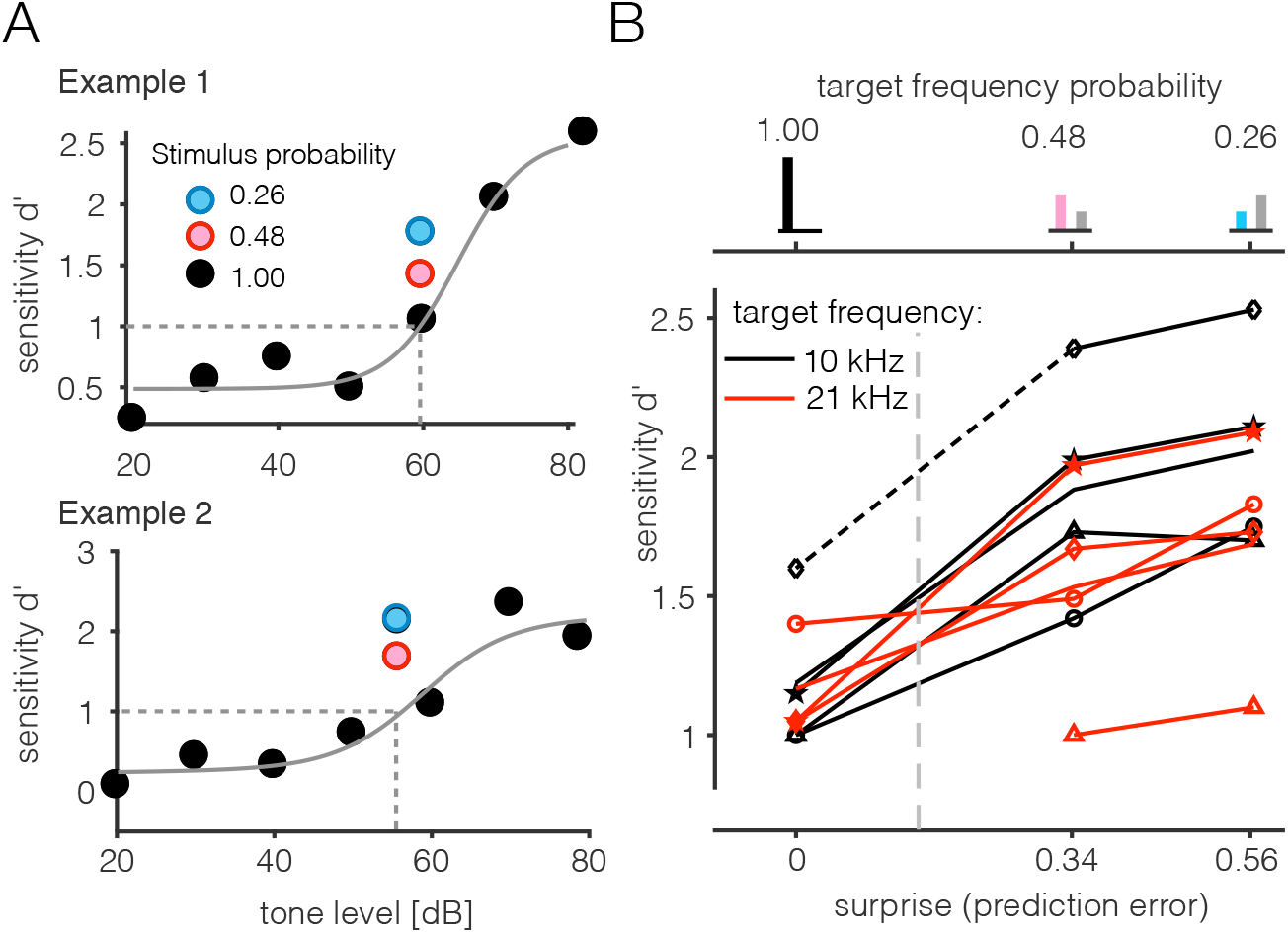
Results for Experiment 1 - tone in noise detecion. (A) Example performance of two different animals for the tone-in-noise stimuli at a single tone frequency, presented with different probabilities. Before the mixed-frequency experiments, animals were tested individually for their thresholds at each tone frequency by presenting tones of a single frequency (probability 100%) at different levels to construct psychometrics functions (black circles, grey line). In the mixed experiments, tones with a level corresponding to a d’ of 1 (dashed line) were presented with probabilities of 48% (red circle) or 26% (blue circle). (B) Population data for all four animals at the two different frequencies used (red: 21kHz, black: 10kHz). The values for a probability of 100% were taken from the psychometric function obtained after the mixed experiments. Histograms above the graph visualize the probability of the tone in the respective sessions, the x-axis shows the surprise quantified as the prediction error. Note that larger numbers indicate more surprising stimuli. Total number of sessions included: 208 (152 for the mixed session, 56 for the psychometric functions).

### Experiment 2: Frequency change detection in streams

Contrary to the behavior displayed in experiment 1, a strategy to focus on high-probability sounds would have maximized rewards. A possible explanation for mice not taking advantage of tracking probabilities is that they are not able to focus on a single frequency band in a continuous noise background with very sparse tones appearing at random times. We reasoned that a more natural situation could be the presence of multiple streams of tones that allow to attach selective attention to one of these streams (Lakatos et al., 2013; Schwartz and David, 2018). We therefore designed an experiment in which animals had to detect a frequency change in either one of two continuous streams of tone pips (Fig. 3A). The repetition rate was rapid (5 Hz for either stream) and the tone streams were more than an octave apart in frequency, a parameter range that results in a clear two-stream percept in most animals, including rodents (Itatani and Klump, 2017; Noda et al., 2013). To this end, we trained a new batch of naïve mice (n=6). Again, we varied the probability that a target could appear in either of the two streams. In one set of sessions, frequency changes would be inserted in either one of the two streams only. In a second set, targets appeared in both streams with equal probability. Sessions were randomized in order to avoid sequence effects.

**Figure 3.**
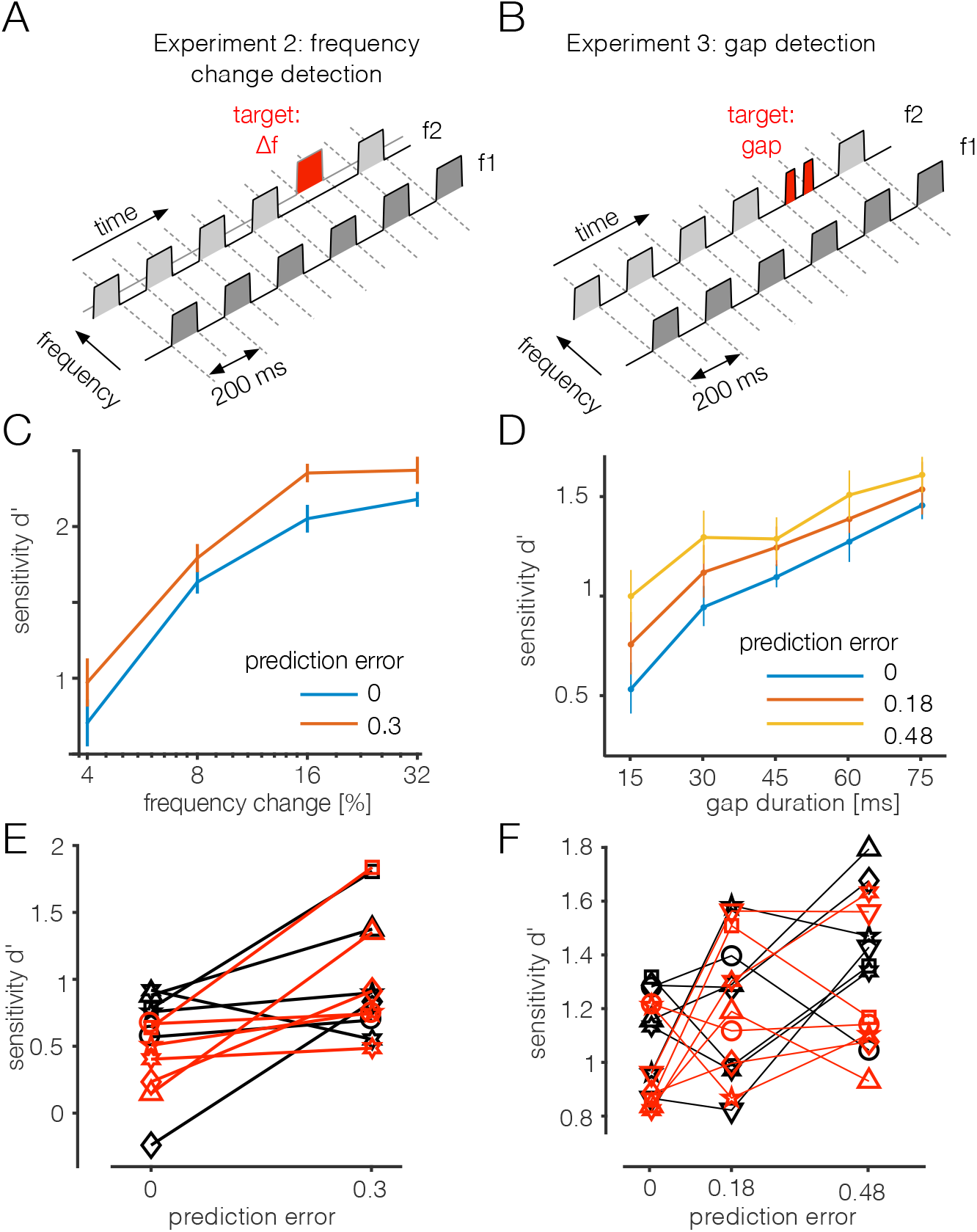
Stimulus paradigms and results for Experiments 2 and 3. (A) Paradigm for Experiment 2: Two continuous, interleaved streams of tone pips with different frequencies were presented. f1 = 10kHz, f2 = 21kHz. Animals had to detect a change of the frequency in either of the two streams. (B) Paradigm for Experiment 3: A short gap was inserted into one of the the two narrowband noise streams (center frequency same as f1 and f2 in A) as a target for detection. (C) Mean performance of all animals (n=6) in Experiment 2 tested at different values of frequency change at either 50% (red) or 100% (blue) probability. Errorbars show standard error of the mean (SEM). (D) Mean performance of all animals (n=7) in Experiment 3 tested at different gap durations at 33.3% (yellow), 66.7% (red) or 100% (blue) probability. Errorbars deptic SEM. (E) Sensitivity as a function of prediction error for Experiment 2 - tone change detection - for all animals tested in both frequency streams (black: 10 kHz, red: 21kHz). Each line joins data from an individual mouse for targets with a frequency change of 16%. (F) Sensitivity as a function of prediction error for Experiment 3 - gap detection - for all animals, mean across all gap durations in either of the two frequency streams (black: 10kHz, red: 21kHz).

When we compared the mean sensitivity for the two different probability levels, we observed a higher mean sensitivity for the mixed sessions for all tested frequency changes (Fig. 3C). As in Experiment 1 (Fig. 2), targets were more salient to the mice if they were distributed between the two streams than if they were played in one of the two streams only. This was confirmed when we compared all animals for both streams (Fig. 3E; rmANOVA, F = 6.0, p=0.0171). Experiment 2 confirmed that the animals are able to track probabilities from session to session, but detectability is determined by surprise rather than by expectation-driven attention, despite the latter being the better strategy to maximize rewards in a given session.

### Experiment 3: Gap detection in streams

Since the two streams were separated by frequency and target changes were along the same dimension, we aimed to test whether our results would generalize to other stimulus dimensions. Therefore, we trained a new set of animals (n=7) to detect temporal irregularities in the form of short gaps introduced into one of the two streams (Fig. 3B). Here, we used three probabilities for each condition: targets in only one of the two streams (100%), or 66.7% and 33.3% probability in sessions with targets in both streams. As already observed for the frequency changes, sensitivity for detection of gaps strongly depended on target probability, with the best detectability for low probability targets in the mixed sessions, and lowest detection performance for targets in only one out of two streams (Fig. 3D). We observed this effect for both possible target streams in all animals (Fig. 3F, rmANOVA, F(1,206) = 30.0, p<0.001). Experiment 3 confirmed our results from the previous experiments and generalizes the saliency of surprising targets to temporal features as well.

### Probabilistic choice model

We observed higher detection performance for low-probability stimuli in three different behavioral experiments. However, this does not necessarily mean that the animals were tracking long-term probability. When manipulating probability, the structure of the randomized trial sequences is changed as well: In sessions in which one type of target is presented with low probability, stimuli are more often preceded by a different target than if presented in high-probability sessions. A simple attentional switch after each trial could explain our results just as well as tracking probability over a time course of minutes to hours. In order to test whether the animals were tracking probability over longer time-scales or simply displaying short-term trial-history effects, we devised a probabilistic choice model (Fig. 4A). The model included the factors stimulus intensity, stimulus probability within the session, and recent history of stimuli presented in the immediately preceding trials. The model was fit separately for each mouse and experiment, in versions including or excluding probability and history terms. If the probability-dependence was due to recent history effects, a model including only the respective term should perform equally well as one including both probability and history, and better than one that takes only probability into account. Inclusion of the probability term significantly improved model performance (Fig. 4B). In contrast, inclusion of the recent-history term (up to four preceding trials) improved the model only marginally (Fig. 4B).

**Figure 4.**
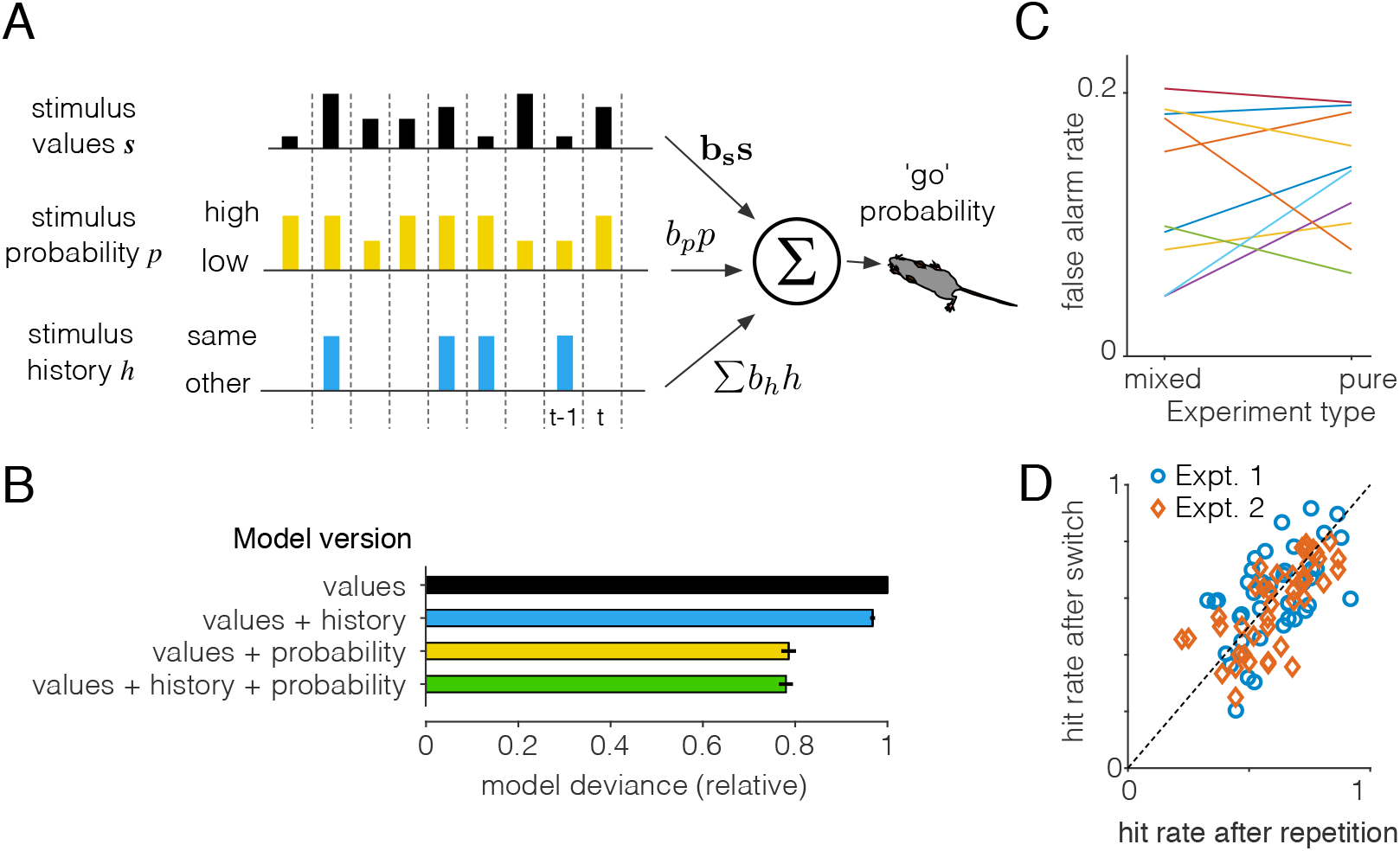
Probabalistic choice model. (A) Schematic illustration of the probabilistic choice model. The full model includes stimulus intensity, stimulus probability within the session for each stimulus, and recent history of stimuli presented in the immediately preceding trial steps *t-i*. Stimulus values were different in each experiment: signal-to-noise ratio in Experiment 1, frequency shift in Experiment 2 and gap duration in Experiment 3. The model was fit for each mouse and experiment, in four different versions, including either all three factors, stimulus values only, values + history, or values + probability. (B) Performance of the four model versions, plotted as deviance of model output to the data, relative to the model including stimulus values only. Note that smaller numbers mean better model performance. Bars represent mean deviances from all animals in the three experiments ±SEM (C) False alarm rate depending on whether stimuli of one class were presented as the only stimuli in the session (‘pure’) or whether they were combined with other stimuli (‘mixed’). Each line represents mean false alarm rates from a single animal. (D) Influence of immediate trial history on hit rate. x-axis: hit rate when the stimulus in the trial before was drawn from the same class as the current stimulus (‘repetition’), y-axis: hit rate to stimuli that were preceded by a stimulus from another class (‘switch’).

The average interval between two trials was 30.2±10.3s (mean ± standard deviation, n=528 sessions from all three experiments). Since there was little effect of recent trial history up to at least 4 trials, perception in the mice was apparently shaped by long term probability on the time-scale of several minutes at least. In line with this, we could not find a difference between hit rates after a switch of the stimulus class between two trials or a repetition of stimuli from the same class (Fig. 4D). We also did not find a change of overall strategy between mixed and pure sessions - false alarm rates did not differ between those session (Fig. 4C).

## Discussion

Stimulus statistics in auditory scenes have been suggested to shape auditory perception in two contrary ways: (1) A focus on novelty detection, favoring low-probability sounds (Khouri and Nelken, 2015) and (2) focusing attention on expected, high-probability sounds, thereby maximizing overall detection rate (Scharf et al., 1987; Wolmetz and Elhilali, 2016). Here we tested whether attention in mice is drawn rather towards low- or high-probability target sounds. To this end, we conducted three different experiments varying the probability of targets. While humans direct their attention to the most probable target out of several acoustic channels or streams, target detectability in mice decreased with increasing probability. Thus, the more surprising a stimulus was, the more reliably it was detected. This was confirmed in three independent experiments using three separate sets of animals, one with changing probability of target frequency in noise (Fig. 2) and two using a streaming paradigm (Fig. 3) with either a spectral or temporal variation to be detected. Finally, our probabilistic choice model best predicted animal behavior for all three tasks if it took overall probability into account, but not if we considered recent trial history (Fig. 4B). These results suggest that mice indeed track probability over a time scale of at least several minutes, but do not use this information in the same way as humans: instead of maximizing reward by focusing on high-probability targets, the saliency of a target is determined by surprise.

### Different strategy or mouse-specific auditory processing?

It seems that mice are very good at something that humans find hard and vice versa. Are our results in mice really caused by a different strategy with respect to target probability or can it be explained by more basic differences in their auditory system? Mice have much wider auditory filters (Lina and Lauer, 2013), so our stimuli could have been merged into one perceptual category, such that no separate streams would have built up. However, in all three experiments, we used targets that were more than an octave apart, far above the frequency discrimination threshold of mice (de Hoz and Nelken, 2014). It was not assessed whether the two sequences used in experiments 2 and 3 resulted in a streaming percept, as this was not the focus of our study and our results do not critically depend on the sequences being perceived as streams. However, all animal species that have been tested so far showed evidence for streaming for stimuli separated by one octave and upwards (Itatani and Klump, 2017), including rats (Noda et al., 2013), which have similar auditory filters to mice. Furthermore, our positive results on the effect of target probability and its generalization across paradigms provide evidence for perceptual separation rather than merging.

### Deviance detection in the auditory system

We found clear evidence for mice to favor unexpected, surprising stimuli. This finding suggests that saliency of a target is determined by it deviating from previous acquired prior probabilities. An explanation for this behavior is that in mice other than humans, the deviance signal (prediction error) is not weighted by probability-cued attention. There is a large body of work on enhanced neural representation of deviant stimuli in the auditory system in both animal models and humans. At the single cell level, stimulus specific adaptation (SSA) describes the enhancement of the neural representation of low-probability sounds (Khouri and Nelken, 2015; Carbajal and Malmierca, 2018). A typical paradigm is the presentation of a sequence of tones of two different frequencies, with varying relative probability (Ulanovsky et al., 2003). Experiment 1 of this study is such a paradigm, but the ratio of stimulus duration (0.5s) and the very long inter-stimulus interval (mean of all sessions: 30.2s) has not been reported before. However, time scales up to several minutes are reasonable based on the measurement of adaptation time constants in cat auditory cortex (Ulanovsky et al., 2004). The stimuli in our Experiments 2 and 3 extended this to two synchronously presented sequences of standard and deviants - Experiment 2 using frequency shifts and Experiment 2 using temporal deviants. SSA is likely to shape responses to the deviant targets in both of the streams in either experiment. Deviants in both streams are very rare: ~30s inter-trial intervals with a background pulse repetition rate of 5 Hz result in a deviant probability of ~1% in session with target in one stream and ~0.5% in sessions with equal distribution between the two streams. Different performance for these sessions would imply that SSA has to reflect differences in probability as small as Δ0.5%. Neural sensitivity for such small changes has not been reported yet, but there is no principle reason why they cannot exist. Alternatively, SSA could work on a higher structural level, reflecting the task and the auditory scene as a whole. SSA has been shown to extend beyond simple pure tone patterns (Nelken et al., 2013) and to more complex statistical structure of the sensory context (Yaron et al., 2012). Similar to the findings presented here (Fig. 4), SSA is sensitive to average statistics rather than recent history (Rubin et al., 2016). Not only in animal models, but also in humans, deviance detection represents one of the main principles of auditory processing. It is reflected by the mismatch negativity (MMN) component of the EEG and is present for a large range of stimuli and time scales (Näätänen et al., 2007, 1978). Both SSA and MMN are discussed as a response to the violation of expectation (Heilbron and Chait, 2017; Khouri and Nelken, 2015; Carbajal and Malmierca, 2018) within the framework of predictive coding (Friston, 2005).

Despite the pervasive presence of neural signatures of deviance detection in auditory systems, it is very difficult to directly observe a correlate at the behavioral level. This is probably due to its pre-attentive nature - deviance detection is observable in passively listening subjects (Tiitinen et al., 1994) as well as in anaesthetized animals (Parras et al., 2017). In active listening tasks, implicit cueing could then use the predictive signal to channel selective attention to high-probability sounds (Wolmetz and Elhilali, 2016) by inverting the sign (Kok et al., 2012). However, in mice we may be able to directly observe the strength of a prediction error signal at the perceptual level.

### Sensory ecology

Deviance detection is of foremost importance for the detection of sudden, potentially dangerous changes in the environment. In mice, as potential targets of predation, deviance detection may have priority over probability-cued selective attention. Contrary, both humans and carnivores (Schwartz and David, 2018) may be cued implicitly using target probability - providing evidence that the effect of stimulus probability on perception may be due to sensory ecology rather than taxonomy. Both primates and carnivores might use their auditory senses to tune in and follow potential prey or conspecific communication signals. Interestingly and consistent with the strategy reported her for mice, threatening stimuli in humans are best perceived if they occur with relatively low-probability (McFadyen et al., 2019). In contrast, implicit cueing in reward-based tasks usually enhances perception of high-probability signals (Girshick et al., 2011; Scharf et al., 1987; Wolmetz and Elhilali, 2016).

An alternative explanation for our results would be that mice are not able to provide top-down influence and scale the deviance signals accordingly. However, recent work suggests that mice are able to selectively attend at least *explicitly* cued visual patterns (Wang and Krauzlis, 2018) or auditory streams (Chapuis and Chadderton, 2018). This could indicate that mice do not lack a mechanism for top-down attentional control of input signal scaling, but it is not activated by probability-cueing. Instead, if mice use contextual auditory information mainly for the detection of threats, this rule may be hard-wired and not under the control of top-down signals.

Our results suggest that in mice, predictions based on complex statistic regularities are computed along the sensory pathway, but mostly used to suppress ongoing input, similar to sensory adaptation on shorter time scales (Carbajal and Malmierca, 2018; Parras et al., 2017). The development of attentional modulation of error signals in carnivores and primates may have been added later on to this first step of probabilistic analysis of complex sensory scenes.

### Perspective

In summary, our study provides the first evidence for animal detection behavior being shaped directly by prediction error. This finding could be very helpful for future work on prediction-guided behavior, since we may be able to study the neural mechanisms underlying extraction of complex contextual sensory information without the confounding of the interplay with (top-down) attentional modulation shaped by the task. The mouse model offers unrivaled possibilities to record and manipulate neural activity in the behaving animal. In future studies this may not only enable measurements of neural deviance detection during relevant behavior. It also offers the perspective of direct manipulation of potential mechanisms, with the observed behavior as readout to infer causal relationships.

## Methods

### Animals

In total 17 adult male mice bred at the University of Oldenburg animal facilities were used in the experiments (Experiment 1 - n = 4, Experiment 2 - n=6, Experiment 3 - n=7). All mice had a C57BL/6.CAST-*Cdh23^Ahl+^* background (the Jackson laboratory, #002756) and were between 3 and 9 month old. We used this line because it does not display the age-dependent hearing loss which is present in other C57BL/6 lines (Johnson et al., 1997; Kane et al., 2012). Animals were kept at a reversed 12/12 hour dark-light cycle, all experiments were performed during the dark period. Animals had unlimited access to water but were food-deprived to a moderate extent (85-90% of their *ad libitum* weight) and single-housed in standardized cages but with visual and olfactory contact to neighboring animals. Cages were equipped with cage enrichment. All experiments were approved by the responsible authorities (Lower Saxony State Office for Consumer Protection and Food Safety, license number 33.9-42502-04-13/1271).

### Behavioral paradigm

All three experiments were performed using the following reward-based go/no-go paradigm. Animals were placed one an annular platform made from wire mesh (Fig. 1A). The raised platform was placed in a custom sound-proof chamber that was lined with pyramid foam. On one side of the platform, a small pedestal was installed. Once the animals ascended the pedestal, a random, variable waiting time started, ranging from 1.25 to 5.25s. After this random interval, a target was presented.

The onset of the target triggered a 1s response window. If the animals descended within the window (,go’), a food pellet (0.02g, Dustless precision pellets rodent, grain based, Bio-Serv, #F0163) was delivered at the opposite side of the annular platform. If the animals stayed on the pedestal, a new trial was presented after a newly drawn waiting time. In order to estimate how many of hits were awarded by chance, about one third of trials (depending on experiment) were unrewarded sham trials, with the same distribution of waiting times as the target trails. Neither false alarms nor misses were punished or rewarded. A typical session contained 60 targeted trials and 25 sham trials and lasted 30-40 minutes. Animals were tested once per day. All experiments were controlled by custom Software (Github Repository: https://github.com/Spunc/PsychDetect) written in MATLAB (The Mathworks). Pellet dispenser and light barriers were custom build (University of Oldenburg workshop) and controlled by a microcontroller (Arduino UNO, Arduino AG, Italy) connected to a Windows PC.

### Stimuli

For sound presentation a speaker (Vifa XT 300/K4, Denmark) was mounted in the sound-proof chamber approximately 0.5m above the pedestal. Sound was generated using a high-fidelity sound card (Fireface UC, RME, Germany) connected to the PC. Sound was played back at either 192kHz (experiment 1) or 96kHz (experiments 2 & 3) sampling rate. The speaker was calibrated at the approximate position of the head of the animals using a measurement microphone (model 40BF, G.R.A.S, Denmark).

#### Experiment 1 - Tone in noise detection

Tones in noise served as a target in Experiment 1. Once a session started, broad-band noise (4-64kHz, 60dB) was constantly played until the end of the session. Pure tone of either 10 or 21kHz served as targets (2ms cosine ramps, 500ms duration). In the sessions containing only one target frequency, the level for that frequency was varied between 20 and 80dB in steps of 10dB in order to obtain a psychometric function. Psychometric functions were fit with a logistic function and an individual signal-to noise ratio (SNR) threshold was estimated. In the mixed sessions, we used the level corresponding to the individual SNR thresholds, estimated as the point on the psychometric curve with a d’ value of 1. During the mixed session, the first 10 trials were taken from either of the two frequencies (priming frequency). For the rest of the sessions, both frequencies were played back with equal probability. In addition, target tones of a third frequency close to the priming frequency were played with equal probability. These stimuli were not used for further analysis. Only the later part of the session was used for analysis of the animal’s performance. Each animal performed at least 10 session for both priming frequencies. Measurement of psychometric function was repeated after the mixed sessions in order to rule out effects of perceptual learning when comparing single-frequency with mixed sessions.

Prior to the described experiments, animals have been trained in a tone in noise detection task with either one of the two target frequencies (10 or 21kHz) which were randomly chosen for each session until the performance reached a stable level in several consecutive sessions.

#### Experiment 2 - Frequency change detection in streams

For Experiment 2, two alternating tones with frequencies of 10 and 21kHz (1.07 octaves) were played at rate of 5 tones/s throughout the experimental session. Tone duration was 100ms including 2ms cosine ramps. The level of each individual tone was roved between 60 and 66dB SPL (randomly) in order to avoid the detection of a differences in loudness when the shift in frequency occurred. The frequency of a tone from either tone sequence was shifted upwards by 4%,8%,16% or 32%. Mice had to report the appearance of the frequency shift within 700ms after onset of the shifted tone. Within a session, targets appeared either in only one of the two tone sequences (,single’) or with 0.5 probability in either of the two sequences (,mixed’). Each animal completed at least 8 sessions for each of the mixed session types.

Prior to experiments, animals have been trained in a frequency change detection task in a single stream, frequencies (10 or 21kHz) were chosen randomly for each training session. After receiving a stable and similar performance in both streams in several consecutive sessions, experiments with the two alternating streams were introduced. Mice responded towards the simultaneous presentation immediately without a decline in performance.

#### Experiment 3 - Gap detection in streams

The temporal structure of the sequences in Experiment 3 was the same as in Experiment 2, but instead of pure tones, narrowband noise with a bandwidth of 0.25 octaves around 10 or 21kHz was used. We introduced this adjustment, because mice were not able to detect temporal gaps in the tone streams used in Experiment 2. The level of narrow band pulses was fixed at 60dB SPL. In the target pulses, gaps with duration of 15, 30, 45, 60 and 75ms were introduced (including 2ms cosine pulses). The response window was 1 s. For Experiment 3, we used three different probabilities: 1 (target only in one sequence), 0.66 or 0.33. Each animal completed at least 8 sessions for each session type.

For this specific paradigm, the auditory training was performed in the same way as experiment 2 but was started with a broadband instead of a narrowband noise, center frequencies of 10 and 21kHz were randomly chosen. After animals showed a stable performance for both individual streams, the bandwidth of the noise was slowly reduced in each session until the final bandwidth of 0.25 octaves was reached. Subsequently, final experiments with both streams were conducted.

### Data analysis and statistics

In all three experiments, for each session *i* and stimulus class *s*, the sensitivity *d′* was calculated as:

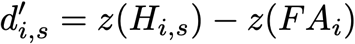

where *z()* is the inverse of normal cumulative function, *H_i,s_* is the hit rate for the stimuli with parameters *s* in the ith session *P(response|stimulus* s) and *FA_i_* is the false alarm rate *P(response|sham)*.

In order to check for significant effects of stimulus probability on the sensitivity, we fit a generalized mixed effects model (MATLAB *fitglme*), with the d’ values as response variable and probability and stimulus parameters as factors. In Experiment 1, the stimulus parameter factor was target tone frequency. For Experiment 2, relative frequency shifts were entered as factor. For Experiment 3, the stimulus factor was gap duration. For each experiment, we performed repeated measures ANOVA (rmANOVA, MATLAB) and report both F-values and exact p-values up to the fourth decimal.

### Probabilistic choice model

To account for different factors affecting animal choice behavior we devised a probabilistic choice model, similar to what has been used before in order to include history in psychophysics (Busse et al., 2011).

The probability *p_go_* to jump at a given trial *t* in a behavioral session is given by:

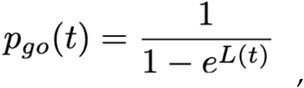

with the response variable *L(t)*, that is a weighted sum of three main terms: (1) the stimulus parameters ***s**(t)*, (2) the overall probability of the stimulus to appear in the given channel *p(t)*, and the stimulus history ***h**(t)*:

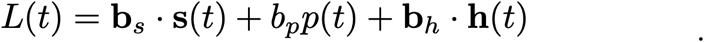

The stimulus parameters depend on the paradigm. For Experiment 1, this was absolute stimulus frequency and the signal to noise ratio. For Experiment 2, this was the absolute frequency of the stream the target appeared in and the frequency shift of the target. For Experiment 3, we entered absolute frequency of the target stream and the gap duration in the target pulse.

The probability term is constant across a given session and only depends on the target channel. The history term is described by:

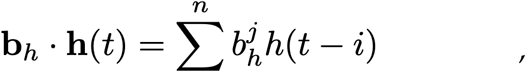

where *h(t-i)* is is 1 if the target in the *(t-i)*th trial before the current was in the same channel and 0 if it was presented in the respective other.

The weights were fit using the Matlab function *glmfit* with a logit link and no constant term. For each animal, sessions were combined into sets that each contained all probability distributions (four single sessions in Experiments 1 and 3, three sessions in Experiment 2). For each experiment and animal, at least five such sets were combined randomly and corresponding models were fitted, resulting in a total of 86 sets. For each such set four versions of the model were fitted, the full model above and the following reduced versions.

Stimulus parameters only:

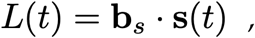

stimulus parameters + probability,

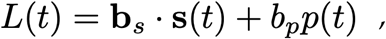

stimulus parameters + history:

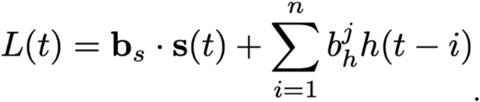

For each set and model version, the deviance between the animal’s response and the probability *p_go_* was collected and normalized to the model deviance for the model version including stimulus parameters only.

## Acknowledgements

The authors thank C Koeppl, MS Malmierca, and GV Carbajal for their helpful comments and productive scientific discussions and D Gonschorek for the excellent support. Additionally, the authors would like to thank M Maleki and AA Korte for the support with animal handling and behavioral training.

